# Interleukin-27-induced HIV-resistant dendritic cells suppress reveres transcription following virus entry in an SPTBN1, Autophagy, and YB-1 independent manner

**DOI:** 10.1101/2023.06.12.544550

**Authors:** Tomozumi Imamichi, Qian Chen, Bharatwaj Sowrirajan, Jun Yang, Sylvain Laverdure, Anthony R. Mele, Catherine Watkins, Joseph W. Adelsberger, Jeanette Higgins, Hongyan Sui

**Author notes:** These authors contributed equally to this work (Tomozumi Imamichi, Qian Chen, Bharatwaj Sowrirajan).

## Abstract

Interleukin (IL)-27, a member of the IL-12 family of cytokines, induces human immunodeficiency virus (HIV)-resistant monocyte-derived macrophages and T cells. This resistance is mediated via the downregulation of spectrin beta, non-erythrocytic 1 (SPTBN1), induction of autophagy, or suppression of the acetylation of Y-box binding protein-1 (YB-1); however, the role of IL-27 administration during the induction of immature monocyte-derived dendritic cells (iDC) is poorly investigated. In the current study, we investigated the function of IL-27-induced iDC (27DC) on HIV infection. 27DC inhibited HIV infection by 95 ± 3 % without significant changes in the expression of CD4, CCR5, and SPTBN1 expression, autophagy induction and acetylation of YB-1 compared to iDC. An HIV proviral DNA copy number assay displayed that 27DC suppressed reverse transcriptase (RT) reaction without influencing the virus entry. A DNA microarray analysis was performed to identify the differentially expressed genes between 27DC and iDC. Compared to iDC, 51 genes were differentially expressed in 27DC, with more than 3-fold changes in four independent donors. Cross-reference analysis with the reported 2,214 HIV regulatory host genes identified nine genes as potential interests: Ankyrin repeat domain 22, Guanylate binding protein (GBP)-1, -2, -4, -5, Stabilin 1, Serpin family G member 1 (SERPING1), Interferon alpha inducible protein 6, and Interferon-induced protein with tetratricopeptide repeats 3. A knock-down study using si-RNA failed to determine a key factor associated with the anti-HIV activity due to the induction of robust amounts of off-target effects. Overexpression of each protein in cells had no impact on HIV infection. Thus, we could not define the mechanism of the anti-HIV effect in 27DC. However, our findings indicated that IL-27 differentiates monocytes into HIV-resistant DC, and the inhibitory mechanism differs from IL-27-induced HIV-resistant macrophages and T cells.

## Introduction

Dendritic cells (DCs) regulate the immune response by initiating and shaping adaptive immune responses against pathogens and cancer. In the periphery, immature DCs (iDC) surveil their environment for foreign pathogens [1, 2]. The uptake of pathogens by iDC in peripheral tissues causes the DCs to migrate to the lymph nodes. During this process, iDC upregulate activation markers and degrade foreign pathogens into peptide antigens [3]. Mature DCs (mDCs) present these peptide antigens on their human leukocyte antigen (HLA) molecules to activate and clonally expand T cells and clear the pathogen [4, 5]. However, certain pathogens, such as human immunodeficiency virus type 1 (HIV-1), can hijack these pathways in DCs to mediate virus replication and dissemination throughout the body [6].

HIV-1 infects DCs via two distinct pathways, *cis*- and *trans-*infection. *Trans*-infection requires the endocytosis of the entire HIV-1 virion into endosomal vesicles, which are subsequently released into the immunological synapse between DCs and T cells via exocytosis [7–9]. The *cis*-infection of DCs by HIV-1 results in productive infection, leading to the generation of new virions [10]. The productive infection of DCs requires HIV-1 to exploit numerous host cell factors for replication and dissemination. In turn, host cells may express cellular restriction factors that inhibit viral replication. Many of these factors are stimulated upon exposure to certain cytokines, such as type I interferons (IFNs) [11–14]. Our previous studies identified interleukin (IL)-27 as a novel inhibitory cytokine of HIV-1 produced by cervical cancer vaccine-treated macrophages [15], which induces multiple IFN-stimulated genes, including host factors, via an IFN-independent mechanism [16, 17].

IL-27, a member of the IL-12 cytokine family (IL-12, IL-23, IL-35, and IL-39) [18], is a heterodimer composed of the IL-27 p28 and Epstein-Barr virus-induced gene 3 (EBI3) subunit [19, 20]. It plays crucial roles in both innate and adaptive immune responses. It binds to the IL-27 receptor (IL27R), which is a heterodimer made of IL27R alpha chain (also known as WSX-1) and the glycoprotein 130 (gp130) [18, 21]. Stimulation of the IL27R by IL-27 triggers downstream signaling through activation of Janus Kinase (JAK) and Signal Transducer and Activator of Transcription (STAT) pathways [22, 23]. The activated STAT-1 and STAT-3 induce many downstream host-cell gene activation including microRNAs and long noncoding RNAs [16, 24–27]. Previous studies have demonstrated that IL-27 has both pro- and anti-inflammatory functions [23]. Naïve T-cells proliferate and differentiate into T helper 1 (Th1) cells in the presence of IL-27 [28]. In contrast, IL-27 activates regulatory T cells and inhibits the generation of Th17 and T follicular cells. Additionally, it stimulates the production of the anti-inflammatory cytokine IL-10 [29]. IL-27-mediated HIV-1 inhibition occurs via the induction of IFN-stimulatory genes (ISGs) in a type I IFN-independent manner [16, 30]. Furthermore, IL-27 regulates the functions of epidermal cells and keratinocytes [31–33], the central nervous system [34, 35], tumor cells [36, 37] and mucosal innate immune responses [38, 39]. Therefore, IL-27 is considered a pleiotropic cytokine. IL-27-mediated function has been reported to be IFN-induction-dependent and independent [40, 41], cell type-dependent manner [41].

We previously assessed the anti-viral activity of IL-27 in terminally differentiated primary monocyte-derived macrophages (MDMs) and iDC [30, 42]. We infected MDMs or iDC with HIV and cultured them in the presence of IL-27 to examine its anti-HIV effects and observed that IL-27 suppressed HIV infection and replication in a dose-dependent manner [15, 42]. To define a role of IL-27 during monocyte differentiation, the cells were stimulated by differentiation inducing factors (macrophage colony-stimulating factor (M-CSF) or human AB serum) in the presence of IL-27. We found that IL-27-induced MDMs resist infection with HIV-1 [24, 42], and demonstrated that the resistance was associated with the down-regulation of spectrin beta, non-erythrocytic 1 (SPTBN1) or the induction of autophagy [42].

IL-27-induced dendric cells (27DC) are reported to improve antigen processing and stimulate T cells [43], but anti-HIV effect in those cells was unclear. Therefore, in the current study, we examined function of anti-HIV effect in 27DC.

## Material and Methods

### Ethics Statement

Approval for this study, including all sample materials and protocols, was granted by the National Institute of Allergy and Infectious Diseases (NIAID) Institutional Review Board, and participants were provided the informed written consent prior to blood being drawn. All experimental procedures in these studies were approved by the National Cancer Institute at Frederick and Frederick National Laboratory for Cancer Research (the protocol code number: 16–19, approval date: 6 January 2017).

### Cells and reagents

Peripheral blood mononuclear cells (PBMCs) were isolated from healthy donors’ apheresis packs (NIH blood bank) using a lymphocyte separation medium (ICN Biomedical, Aurora, OH, USA), CD14+ monocytes were isolated from PBMC utilizing CD14 microbeads (Miltenyi Biotec, Auburn, CA, USA) as previously described [15]. The purity of the cell types was at least 90%, based on the flow cytometric analysis. Cell viability was determined using the trypan blue (Thermo Fisher Scientific, Waltham, MA, USA). The iDC were generated by culturing CD14+ monocytes in RPMI 1640 (Thermo Fisher Scientific) supplemented with 10% (v/v) fetal bovine serum (FBS) (HyClone/Cytiva, Marlborough, MA, USA), 25 mM HEPES (Quality Biological, Gaithersburg, MD, USA), and 5 µg/ml gentamycin (Thermo Fisher) with 50 ng/ml of IL-4 (R&D systems, Minneapolis, MN, USA) and 50 ng/ml of GM-CSF (R&D Systems) (G4 Media) [30]. To create 27DC, recombinant IL-27 (R&D systems) was added to the G4 media during dendritic cell differentiation at a concentration of 100 ng/ml. Half of the culture media was changed on day 4 with fresh G4 media with or without IL-27 [30]. HEK293T cells were obtained from ATCC (Manassas, VA, USA) and maintained in complete D-MEM (Thermo Fisher Scientific) supplemented with 10 mM HEPES, 10% FBS, 50 μg/mL gentamicin (D10 medium) as previously described [44]. Recombinant SERPING1 was purchased from R& D systems.

### Viruses

Recombinant R5 HIV (HIV_AD8_) and X4 HIV (HIV_NL4.3_) were prepared by transfecting plasmid encoding full-length of each HIV gene (pAD8 [45] and pNL4.3 [46]) into HEK293T cells, respectively. A pseudotyped HIV-1 virus (HIVLuc-VSVG) expressing the luciferase gene (HIV_Luc_) was created by transfection of pHIV-1_NL4/3_-Luciferase-ΔEnv and p-VSV-G into HEK293T cells in 10 cm dishes [42, 47, 48]. The HIV-1_NL4/3_-Env_AD.8_-ΔVpr with Vpr-BLAM was produced by transfection of pHIV-1_NL4/3_-Env_AD.8_-ΔVpr and pVpr-BLAM [49, 50] (a kind gift from Dr. Warner Greene) into HEK293T cells. All plasmid DNA transfections were conducted using TransIT-293 (Mirus, Houston, TX, USA) and Opti-MEM I medium (Thermo Fisher Scientific) following a method previously reported [42]. Supernatants were then ultracentrifuged at 100,000 x g for 2 hours at 4°C onto a 20% sucrose in 10 mM HEPES-150 mM NaCl cushion [51]. Pelleted particles were resuspended in D10 medium and the concentration of HIV p24 was quantitated by using an HIV-1 p24 ELISA Kit (Perkin Elmer, Boston, MA, USA) [44, 51]. Infection titer (50% tissue culture infectious dose, TCID_50_) of each virus was determined by an endpoint assay [52].

### HIV-1 Replication assay

HIV-1_AD8_ (R5 tropic) and HIV-1_NL4.3_ (X4 Tropic) viruses were utilized to infect iDC and 27DC. Cells (5×10^6^) were infected at 5000 TCID_50_ / 1×10^6^ cells at 37 °C for 2 hours on a cell-rotator (Miltenyi Biotec). After infection, cells were washed 3 times with RPMI-1640 supplemented with 10% FBS (RP10) and incubated at 0.5×10^6^ cells/ml in 96 well plates in quadruplicate in G4 media. Cells were cultured for 14 days with one-half of the media being replaced every 3-4 days with fresh G4 media without IL-27. HIV-1 p24 released into the supernatants was measured using the HIV-1 p24 ELISA Kit.

### Pseudotyped HIV Infection

DCs (1×10^6^ cells) were infected with HIV_Luc_-VSVG at a concentration of 500 ng p24/mL in RP10 for 2 hours at 37°C on the cell-rotator. After infection, cells were washed with RP10 followed by being cultured for 48 hours at 0.5×10^6^ cells/mL in 96 well plates in triplicate or quadruplicate in G4 media. Luciferase expression in cells was measured by lysing the cells in Passive Lysis Buffer and incubating the lysis supernatant with the Luciferase Assay System (Promega, Madison, WI, USA). Luciferase activity was read on an Enspire Multimode Plate Reader (Perkin Elmer). Cell lysates were also subjected to a BCA assay for protein quantification (Thermo Fisher Scientific).

### HIV-1 Binding Assay

DCs were incubated with HIV-1_AD8_ (5000 TCID50 per 1×10^6^ cells) at 4°C for 90 minutes to allow binding [53], and then washed to remove unbound virus at 4°C with cold PBS. Total RNA was extracted and reverse transcription of viral RNA was performed as previously described [54]. The levels of virus binding were estimated with copy numbers of viral RNA measured with quantitative RT-PCR [42].

### HIV-1 Fusion Assay

HIV-1 fusion to iDC and 27DC was measured following the previously described protocol [50]. Briefly, 3×10^6^ iDC or 27DC were infected with HIV-1_NL4/3_-Env_AD8_-ΔVpr + Vpr-BLAM at a concentration of 500 ng p24/ml. Infection was allowed to proceed for 2 hours at 37°C. To suppress the virus binding TAK779 (Bio-Techne Corp., Minneapolis, MN, USA) was supplemented during the infection. Following infection, cells were washed 3 times, and stained with the fluorescent dye CCF2 (Thermo Fisher Scientific). After staining, the cells were incubated in CO_2_ independent media with 10% FBS, 25 mM HEPES, 5 µg/ml of Gentamycin, and 2.5 mM probenecid. Incubation occurred for 16 hours at room temperature in the dark followed by analysis by flow cytometric analysis.

### HIV-1 Early and Late Reverse Transcriptase (RT) Products

Quantitative analysis of HIV-1 early and late RT products was performed as previously described [55]. Briefly, 3×10^6^ iDC or 27DC were infected with HIV-1_AD.8_ at 37°C. Infected cells were incubated for 24 hours prior to genomic DNA extraction using the QIAmp DNA Mini Kit (Qiagen). HIV-1 early and late RT products copy numbers were quantitated by real time PCR as described above. HIV-1 late RT products were measured using a primer probe against HIV-1 Gag [56]. HIV-1 early products were measured using a primer probe against the RU5 region of the viral cDNA [57]. Copy numbers for both were normalized to RNase P.

### Fluorescence-activated cell sorting (FACS) Analysis

Total 1×10^6^ cells of iDC or 27DC were washed three times with ice-cold Dulbecco’s PBS (Thermo Fisher Scientific) in the presence of BSA, and then blocked using Fc Receptor Blocker (Innovex Biosciences, Richmond, CA, USA) for 30 minutes at room temperature in the dark. Cells were washed twice in 2% BSA (MilliporeSigma, St. Louis, MO, USA) with 0.5% NaN3 (MilliporeSigma) in Dulbecco’s PBS (DPBS-BSA-NaN_3_). Cells were then stained, including paired isotype controls, for 15 minutes at room temperature in the dark. The following antibodies, and their respective fluorochrome, were used: CD4 (Fluorochrome BV421; BioLegend, San Diego, CA, USA; Ca# 317434) and isotype control (Fluorochrome BV421; BioLegend; Ca# 400158), CCR5 (Fluorochrome APC-Cy7; BD Biosciences, Franklin Lakes, NJ, USA; Ca# 557755) and isotype control (Fluorochrome APC-Cy7; BioLegend; Ca# 400128), CXCR4 (Fluorochrome PE; BioLegend; Ca# 306506) and isotype control (Fluorochrome PE; BioLegend; Ca# 400114). The cells were washed twice in DPBS-BSA-NaN_3_ and then run immediately on an LSR Fortessa flow cytometer (BD Biosciences). The results were analyzed using FCS Express version 7 (DeNovo Software, Pasadena, CA, USA).).

### Western Blotting

Cells were lysed in Radioimmunoprecipitation assay (RIPA) buffer (Boston BioProducts, Milford, MA, UAS) supplemented with 5 mm EDTA (Quality Biologicals), 1X protease inhibitor cocktail (MilliporeSigma) and 1X phosphatase inhibitor (Thermo Fisher Scientific) for 15 minutes on ice, then cell debris were removed by centrifugation at 14,000 rpm for 10 minutes at 4°C. Protein concentration was quantified by using a BCA Assay kit (Thermo Fisher Scientific). A total of 20 µg protein was loaded per sample onto a lane of NuPAGE 4-12% Bis-Tris Gel (Thermo Fisher Scientific) in MOPS buffer (Thermo Fisher Scientific) (for detection of ANKRD22 or SERPING1) or 3-8% TA gels (Thermo Fisher Scientific) in TA buffer (Thermo Fisher Scientific) (for detection of SPTBN1 or STAB1). Gel was subsequently transferred to nitrocellulose membranes and blocked for 1 hour in 5% milk solution in PBS (Quality Biologicals) with 0.1% Tween 20 (MilliporeSigma) (PBS-T). Membranes were then incubated overnight with appropriate antibody at 4°C with gentle agitation. Antibodies against SPTBN1 (Cat# ab72239), ANKRD22 (Cat# ab11638) and glyceraldehyde-3-phosphate dehydrogenase (GAPDH) (Cat# ab 9485) were obtained from Abcam (Waltham, MA, USA). Anti-Stabin-1 (Cat# AB AB6021) and anti-β-actin (Cat# A5316) were obtained from MilliporeSigma. Antibodies were used according to manufacturer’s protocol. Following primary antibody incubation, membranes were washed 3x in PBS-T and then incubated with appropriate secondary antibody conjugated with horseradish peroxidase (GE Healthcare, Chicago, IL, USA). Chemiluminescence was accomplished using Amersham ECL Prime Western Blotting Detection Reagent (GE Healthcare) and signal was detected on the Azure 300 (Azure Biosystem, Dublin, CA, USA). The intensity of the band was analyzed by NIH Image J (http://rsbweb.nih.gov/ij/).

### Autophagy Assay

Endogenous autophagy induction in iDC and 27DC was compared by autophagosome staining as previously described [24, 27]. Briefly, iDC and 27DC were seeded in 96-well plate at 150 x 10^3^ cells/well and then autophagosomes were stained using the Cyto-ID® reagent (Enzo Life Sciences, Farmingdale, NY, USA) and Hoechst 33342 as a counterstain. Autophagosome (2 × 3 composite) images were taken on a Zeiss Axio Observer.Z1 motorized microscope (Carl Zeiss Microscopy, White Plains, NY, USA) at a 10× magnification. Quantification analysis of each image was performed using the Fiji Image J software (National Institutes of Health, Bethesda, MD, USA). For each channel, a background threshold was set up, to create 8-bit masks. For the green channel, the total stained area was retrieved as a measure of autophagosome staining, while a particle counts following watershed processing of the blue channel gave the corresponding cell number, then resulting values display a total cell for each experimental condition [24].

### Two-dimensional Western Blotting

To detect the post-translational modification of YB-1 protein, two-dimensional (2D) WB was conducted as previously described [58]. Whole cell lysate of iDC and 27DC were prepared using RIPA (Boston BioProducts, Milford, MA, USA) supplemented with 5 mM EDTA (Quality Biological), 1X protease inhibitors (MilliporeSigma), and 1X phosphatase inhibitors cocktail (Thermo Fisher Scientific) at 10×10^6^ cells/mL for 15 minutes on ice. The cell lysate was centrifuged at 15,000 x g for 10 minutes at 4°C to remove cell debris. Protein concentration was determined using a BCA Assay. The total cellular protein (100 μg) was subjected for 2D-WB analysis [58].

### DNA Microarray Analysis

DNA microarray assay was performed using the Affymetrix GeneChips (Affymetrix, Thermo Fisher Scientific). The Affymetrix Human Gene 2.0 ST Array was used. Total cellular RNA was extracted from iDC and 27DC using RNA easy kit (Qiagen, Germantown, MD, USA) and quantitated using the 2100 Bioanalyzer (Agilent, Santa Clara, CA, USA) and Qubit Fluorometric Quantitation (Thermo Fisher Scientific). Terminal labeling and hybridization, array wash, stain, and scan were processed according to the Affymetrix recommended standard protocol [30]. Intensity data were processed and summarized to gene level with Partek Genome Suite (Partek, Chesterfield, MO, USA). Differentially expressed gene candidates were selected for verification with an absolute fold change difference >3.0 with significant differences determined by two-way ANOVA. Functional enrichment analysis was performed using Metascape (the database for annotation, visualization, and integrated discovery) [59].

### Quantitative RT-PCR (qRT-PCR)

Total cellular RNAs from iDC and 27DC were isolated as described above. 1 µg of RNA was used to make cDNA using the TaqMan Reverse Transcription Reagents (Thermo Fisher Scientific). Samples were run on Applied Biosystems GeneAmp PCR system 9700 at 25°C for 10 minutes, 48°C for 30 minutes, and finally for 95°C for 5 minutes. DNA products were used for real time qRT-PCR reaction using the TaqMan 2X Universal PCR Master Mix (Thermo Fisher Scientific) on an iQ5 RT-PCR detection system (BioRad, Hercules, CA, USA).

### ELISA

Amounts of the secreted SPERPING1, CCL3, CCL4 and RANTES in culture supernatants was quantified using Human Serpin G1/C1 Inhibitor Quantikine ELISA Kit, Human CCL3/MIP-1 alpha DuoSet ELISA kit, Human CCL4/MIP-1 beta Quantikine ELISA Kit, and Human CCL5/RANTES Quantikine ELISA Kit, respectively following the manufacturer’s protocol. All kits were obtained from R&D systems. Detection limits for quantification of SERPING1, CCL3, CCL4, and RANTES are 313 pg/ml, 31.2 pg/ml, 15.6 pg/m, 31.2 pg/ml, respectively.

### siRNA Transfection

To introduce siRNAs in DCs or 27DC, siRNA targeting gene of interests was transfected into monocytes before differentiation by reverse transfection using a protocol described by Troegeler et al. [60]. Briefly, 200 nmol of siRNA (Thermo Fisher Scientific) in RNase-free water was diluted in 485 µL of Opti-MEM (Thermo Fisher Scientific) and 15 µl of HiPerFect (Qiagen) transfection reagent was added. Transfection mix was gently mixed by pipetting and then incubated at room temperature for 20 minutes.The lipid-siRNA complexes were added to 10×10^6^ monocytes in 1.5 mL of Macrophage-serum free media (Thermo Fisher Scientific) and incubated for 4 hours at 37°C with a gently rotation. After transfection, the supernatants were removed and then the transfected cells were cultured in RP10 containing 50 ng/ml of GM-CSF at 0.5 x10^6^ cells/ml for 16 hr at 37°C to rest. The half of culture medium was replaced with fresh RP10 containing 50 ng /ml GM-CSF and 100 ng/ml IL-4 with or without 200 ng/ml IL-27. The cells were cultured for seven days with changing medium as described above to induce iDC or 27DC.

### Over expression of ANKRD22

To define the impact of ANKRD22 protein on HIV Infection. HEK293T cells were reverse transfected with plasmid encoding each gene. Briefly, 10 μg of pCMV6 encoding ANKRD22 gene (OriGene, Rockville, MD, USA) was incubated with 30 μL of TransIT-293 in 600 uL of OPT-MEM for 30 minutes at RT, and then cultured with 6×10^6^ 293T cells in 10 mL of D10 media in a 100 mm tissue culture dish (Becton Dickinson, Franklin Lakes, NJ, USA) for 2 days. As a control, empty pCMV6 vector (OriGene) was transfected into the cells. The transfected cells were collected and then expression of ANKRD22 was confirmed by WB. HIV infection was conducted using 2 x10^6^ cells in 500 uL of D10 media. The cells were infected with 100 ng p24 /mL of HIVLuc-VSVG for 2 hours at 37°C on the rotator. After infection, the cells were washed with D10 and then cultured at 50×10^3^ cells/ 96 well plate for 2-days in octuplicate in 200 μL D10 media. The cells were lysed using Bright-Glo (Promega) and Luciferase expression was measured using the Enspire Multimode Plate Reader.

### Statistical Analysis

Statistical analyses were performed using GraphPad Prism 9 software (San Diego, CA, USA). Error bars indicate standard deviation (SD) or standard errors (SE). The Student’s unpaired t-test or one way ANOVA or the Mann-Whitney unpaired-test was used and *p* values lower than 0.05 were considered significant.

## Results and Discussion

### IL-27-differentiated monocyte-derived DCs resist HIV-1 infection

To define the function of IL-27 during monocyte differentiation into DCs and HIV susceptibility, freshly isolated CD14^+^ monocytes were cultured in the absence or presence of 100 ng/ml IL-27 in G4 media as described in the Materials and Methods. iDC and iDC induced in the presence of IL-27 (27DC) were infected with an X4-tropic virus, HIV-1_NL4/3_, or an R5-tropic virus, HIV-1_AD8,_ and HIV replication was assessed 14 days after infection, as previously demonstrated [30]. HIV-1_NL4.3_ and HIV-1_AD8_ replication were reduced by 95 ± 3% (n = 4, p < 0.01) and 91 ± 9% (n = 4, p < 0.01) in 27DC, respectively, compared to iDC (**Figure 1A and 1B**). As the replication activity of the X4 virus was low in iDC, as reported previously [30]; further studies were focused on HIV-1_AD8_ infection.

**Figure 1:**
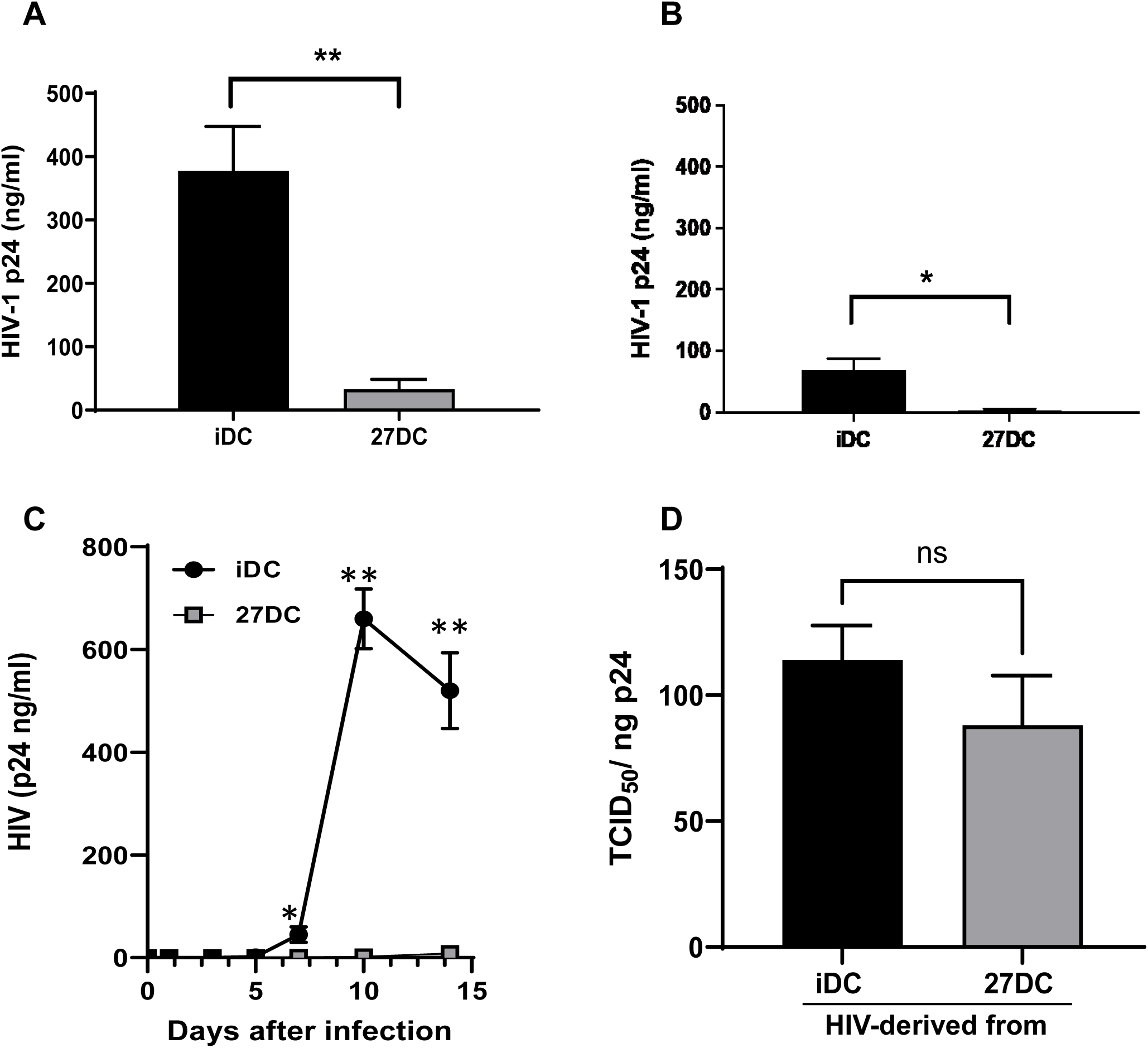
IL-27 treatment during dendritic cell differentiation inhibits HIV-1 replication without affecting infectability. (A and B) iDC and 27DC were infected with HIV_AD8_ (A) or HIV_NL4.3_ (B) at 5000 TCID50/10^6^ cells as described in the Materials and Methods and then cultured for 14 days in G4 media. Viral replication on day 14 was quantified by measuring HIV p24 antigen in the culture supernatants using an HIV p24 antigen ELISA kit. Results are shown means ± SE from four independent donors’ cells. (C) HIV_AD8_ replication kinetics in iDC and 27DC were measured. Aliquots of culture supernatants were collected on Days 1, 3, 5, 7, 10, and 14 after infection. HIV-1 p24 released into the supernatant was measured by ELISA. The result is representative of three different donors; data indicates means ± SD (n=3). (D) HIV infectious titer, 50% Tissue culture infectious dose (TCID_50_/mL) for virus in culture supernatants of HIV_AD8_-infected iDC and 27DC of 14 days post-infection was quantified by the endpoint assay [61] and then normalized with p24 antigen concentration. Results are shown mean ± SE (n=3) of TCID_50_ per ng of p24. Unpaired the Student t test was used to calculate statistical significance, with * representing *p*<0.05, and ** representing *p*<0.01.

To elucidate the mechanism underlying this inhibition, we first measured the HIV replication kinetics over 14 days. In iDC, infection spread was detectable from day 5, and HIV replication was robustly amplified on day 7–10 **(Figure 1C).** In contrast, HIV replication was persistently suppressed in 27DC, with inhibition of replication observed on day 5. This finding indicates that HIV suppression occurs in the early stages of viral replication in 27DC. We then investigated whether the low level of HIV-1 spreading in 27DC was due to the virus particles released from 27DC being less infectious because they contained potential viral-suppressing host factors [62, 63]. Culture supernatants of iDC and 27DC were collected after 14 days of culture and HIV infectious titers in the supernatants were compared. To measure the infectivity of each virus, the 50% tissue culture infective dose (TCID_50_) per ng HIV p24 was quantified using macrophages [54].

Our results from three independent assays indicated that there was no significant difference in the infectivity of virions produced from iDC and 27DC (**Figure 1D**), implying that the inhibition was caused by the suppression of a particular step in the HIV life cycle rather than the incorporation of 27DC-derived antiviral proteins into the virus particles derived from 27DC.

We also analyzed HIV-1’s ability to bind to iDC and 27DC. HIV-1_AD.8_ was incubated with the cells at 4 °C followed by RNA isolation of the virus-cell conjugates. The viral RNA (vRNA) copy number was subsequently quantified in infected iDC and 27DC via qRT-PCR. 27DC expressed 47 ± 10% (*p*< 0.0001, n = 4) less HIV-1 vRNA than iDC **(Figure 2A)**.

**Figure 2:**
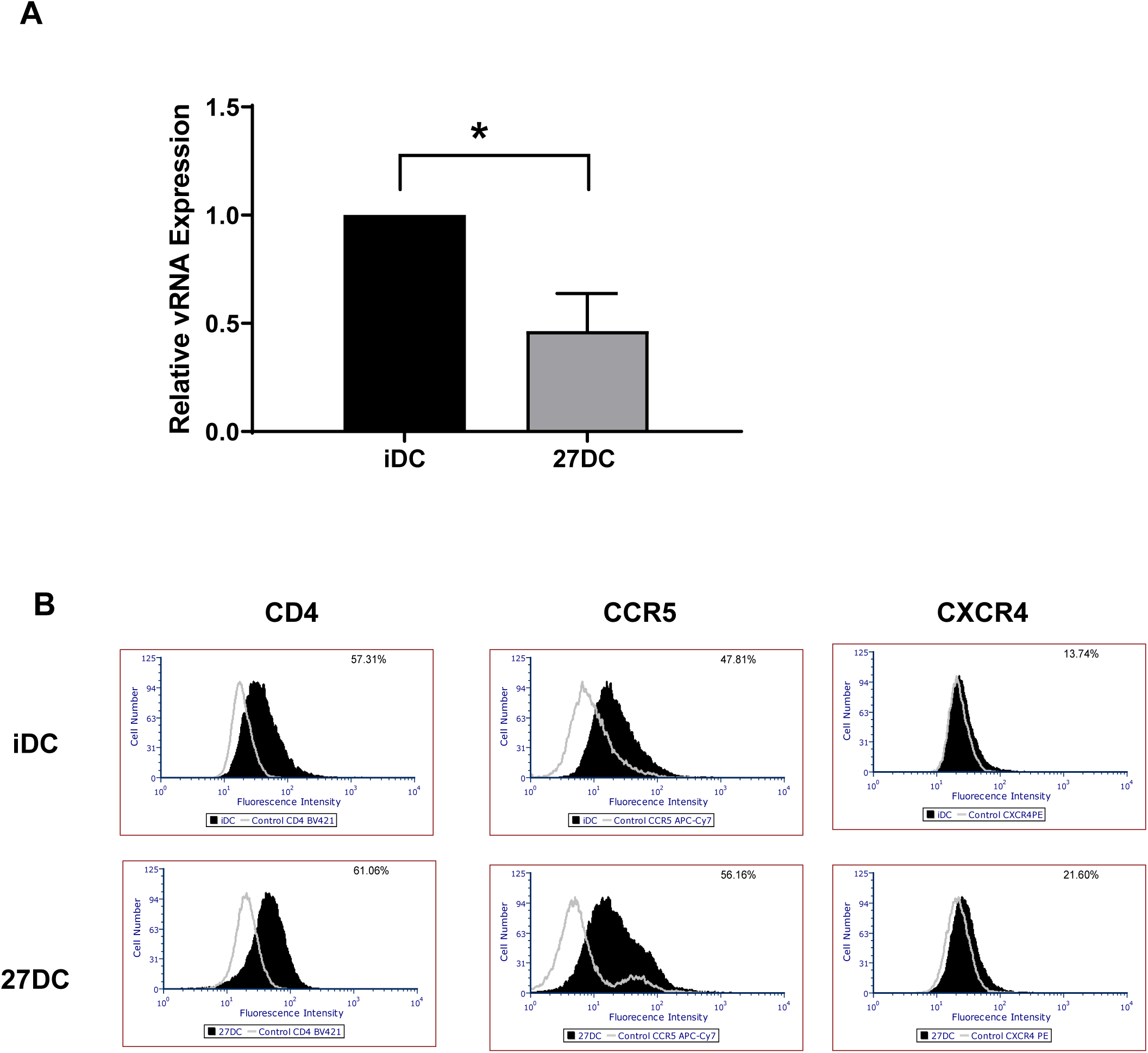
27DC partially resists HIV binding. (A) iDC and 27DC were incubated with HIV-1_AD8_ at 4°C for 90 minutes, and then total RNA was extracted. Bound HIV level and amount of GAPDH RNA were measured using real-time RT-PCR. Copy number of HIV-1 Gag was normalized with GAPDH. The data shown represent means ± SE (n=4). *\* p<0.05,* (B) The expression of CD4, CCR5 and CXCR4 on iDC (top panels) and 27DC (bottom panels) was then assessed by flow cytometry as described in “Materials and Methods.” The staining pattern of isotype control antibodies are shown in open histograms and black histograms represent target protein staining. The x-axis and y-axis show florescence intensity and cell count, respectively. The data are representative of 3 independent experiments with similar outcomes.

The near 50% inhibition of HIV binding might be caused by the downregulation of HIV receptors, CD4 and CCR5. To determine the effect of IL-27 on the expression of CD4 CCR5 on the cell surface, a flow cytometry analysis was performed. CD4 and CCR5 cell surface expression did not significantly change on 27DC compared to that on iDC (**Figure 2B)**. Despite the 50% inhibition of binding, HIV replication in the 27DC was suppressed by more than 90% compared to the iDC. This implies that, in addition to the 50% inhibition of HIV binding, other steps were also influenced in the inhibition. In addition, we also compared CXCR4 expression, 27DC had no impact on the expression (Figure 2).

After the binding of HIV particles to the cell surface, HIV env proteins fuse with the cell membrane, and the core protein in the viral particle is delivered inside the cell. As a result, if the fusion step is further suppressed, an additional inhibition effect between the binding and fusion steps may induce a high level of inhibition in the 27DC. Viral fusion was measured using a Vpr-Blam assay [49]. This assay relies on the incorporation of the β-lactamase-Vpr (Blam-Vpr) chimeric protein into an HIV-1 virion through the co-transfection of pHIVNLAD8ΔVpr and BlaM-Vpr into 293T cells, as described in the Materials and Methods. This virus was subsequently used to infect iDC or 27DC loaded with CCF2, which is a fluorescent dye substrate for the enzyme β-lactamase. Cleavage of the β-lactam ring in CCF2 following infection resulted in a fluorescence change. A CCR5 antagonist, TAK779 was used as a positive control to inhibit binding followed by viral fusion. Un-cleaved and cleaved CCF2 were detected in the AmCyan and Pacific Blue channel, respectively, by flow cytometry. Cleavage of CCF2 indicated the entry of Blam-Vpr into the cell and viral fusion **(Figure 3A)**.

**Figure 3:**
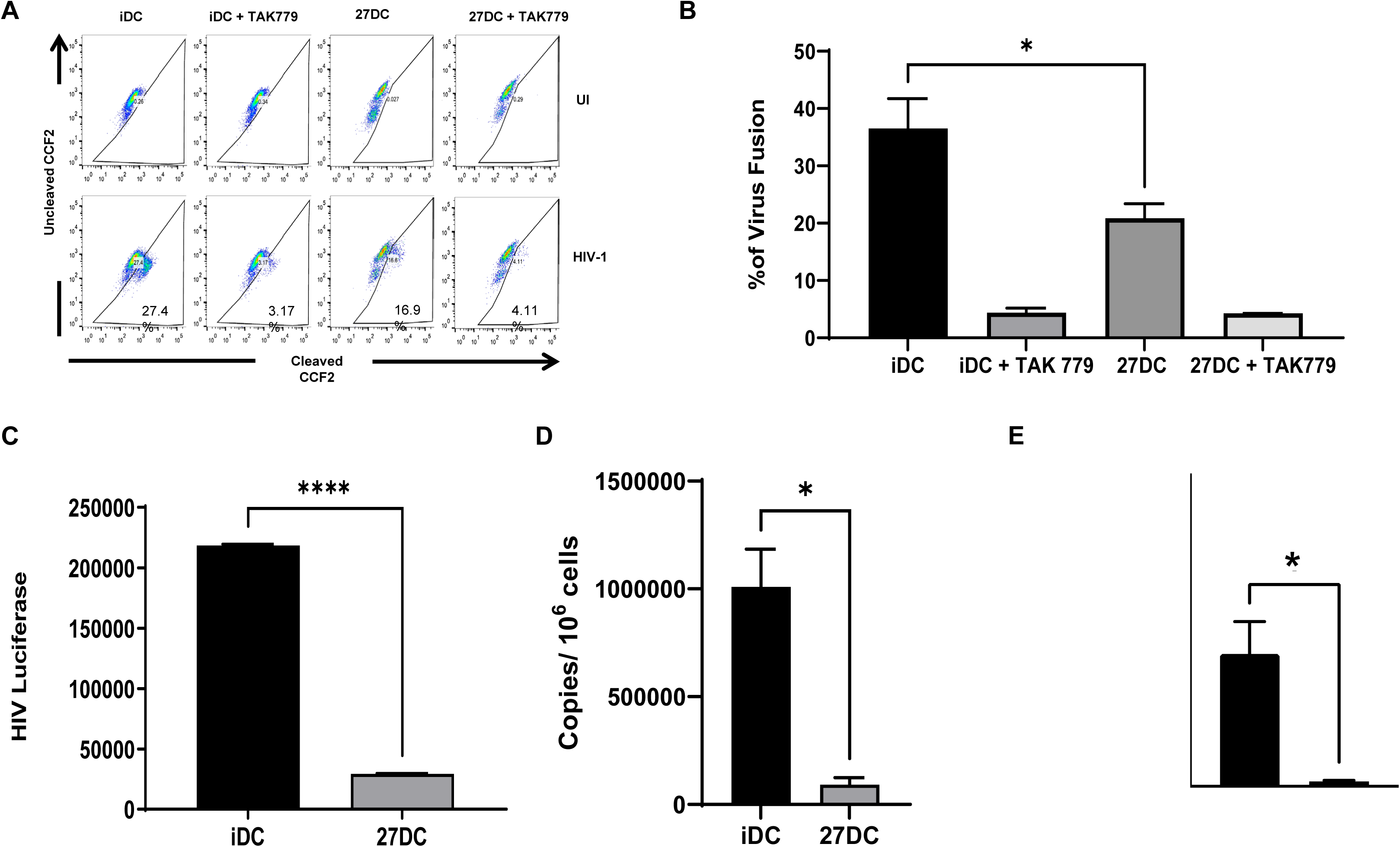
27DC inhibit HIV-1 at both viral entry and before integration. (A) iDC or 27DC were either mock-infected or infected with HIV-1_AD8_ virion containing Vpr-Blam for 2 hours. The cells were loaded with the fluorescent dye CCF2. The cleaved substrate of CCF2-AM by Blam was measured by excitation of the substrate at 410 nm and emission shift from green (520nm) to blue (450 nm) dye. As a positive control for suppression of infection/fusion, cells were treated with 10 nM TAK779 before adding HIV and then infected with the HIV. The dot plot is representative of 4 independent experiments. (B) Percentages of fusion were quantified by detecting CCF2, and the results shown are mean ± SE from four independent studies. (C) iDC and 27DC were infected with pseudotyped HIV-Luc-VSVG for 2 hours and then incubated for 48 hours. Luciferase activity in HIV-Luc-infected iDC and 27DC was measured. The data shows mean ± SD. (D and E) HIV-1_AD8_-infected iDC and 27DC were cultured for 24 hours at 37°C. Genomic DNA was extracted and subjected to quantify proviral DNA copy numbers. Primer/probes against the early RT products (D) and the late RT products (E) were utilized [42]. The result is representative of three different donors; data indicates means ± SD (n=3). **\*:** *p*<0.05, **: *p*<0.01, and ***: *p*<0.001.

Compared to iDC, infection of 27DC with HIV-1 containing Blam-Vpr reduced viral fusion by 43 ± 4% (*p*< 0.05, n = 4) (**Figure 3B**). These findings are consistent with HIV binding inhibition on 27DC described earlier; therefore, it appears that 27DC inhibits HIV infection by approximately 50% in the binding step, without any impact on the virus-membrane fusion step. Although 27DC suppressed HIV at the binding/entry step by nearly 50%, it abolished HIV replication by > 90%, indicating that 27DC further suppressed HIV replication after the fusion step. We previously demonstrated using pseudo-typed replication-incompetent HIV (HIVLuc-VSVG) that IL-27-induced HIV-resistant MDMs suppress the RT step before integration [30]. To elucidate whether 27DC suppressed RT reaction, we used the same technique. HIVLuc-VSVG lacks the HIV envelope gene, encodes a luciferase gene, and expresses a VSVG envelope on the surface of the virus, infecting cells in an HIV receptors-independent manner with a single round of infection and introducing the *Luc* gene. Luciferase activity was therefore considered to correspond to RT activity. The Luc activity in 27DC was found to be nearly 90% lower than that in iDC (**Figure 3C).** HIVLuc-VSVG infection was inhibited in 27DC, thus we presumed that 27DC suppress HIV in a manner like that seen in IL-27-induced HIV-resistant MDMs [42], RT reactions may be suppressed in them. To clarify the suppression, we measured the amount of HIV-1 RT products in iDC and 27DC after HIV infection. The copy numbers of both early and late RT products were measured using qRT-PCR. The 27DC inhibited both late and early RT HIV-1 products compared to iDC (**Figure 3D and 3E).** The early and the late RT product was inhibited by 85 ± 7% and 87 ± 8% (p < 0.05, n = 4), respectively, compared to iDC **(Figure 3D and 3E).** These results indicated that 27DC suppress both HIV-1 binding and RT steps.

### 27DC has no significant impact on SPTBN1 expression, autophagy induction, or post-translational modification of Y-Box 1

We have previously reported that downregulation of SPTBN1 expression [42], induction of autophagy [24], or suppression of acetylation of YB-1 [64] were associated with IL-27-induced HIV-resistant cells; although these changes are cell-type dependent in the IL-27 treatment, these changes are associated with HIV resistance via suppression of the RT step. To determine whether these changes were induced in 27DC and involved in HIV resistance, western blotting and an autophagy assay were performed. SPTBN1 was not detected in 27DC or iDC (**Figure 4A**), and the pI of YB-1 in 27DC was not significantly altered compared to that in iDC (**Figure 4B**).

**Figure 4.**
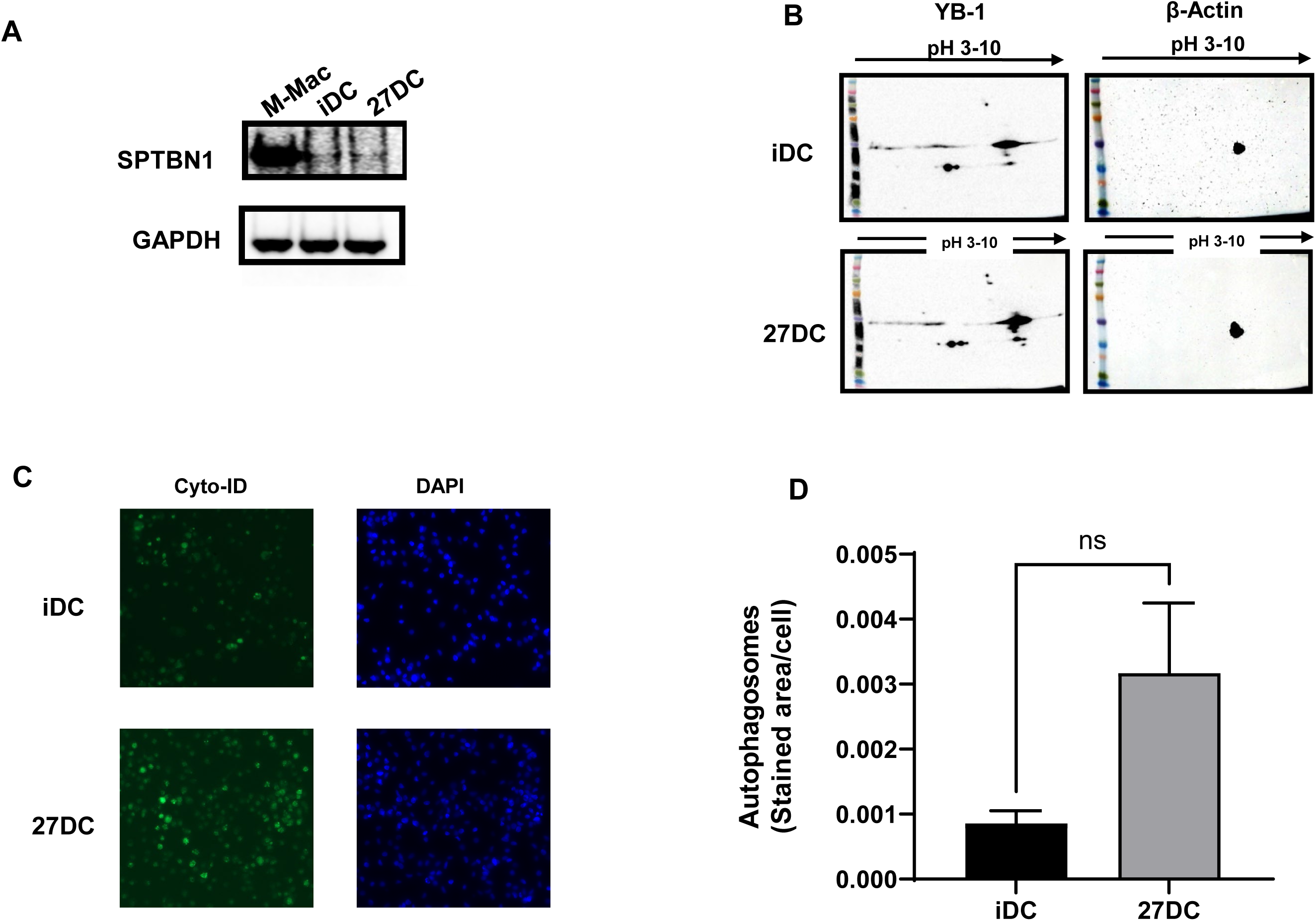
27DC has no impact on the expression of SPTBN1 and PTM of YB1 and the induction of autophagy. (A) Whole cell lysates of iDC and 27DC were subjected for detecting SPTBN1 expression by Western blotting using an anti-SPTBN1 antibody. As an internal control, GAPDH was detected, and as a positive control for SPTBN1 expression, whole cell lysate of macrophage induced by M-CSF was used [42]. (B) The post-translational modification (PTM) of YB-1 was measured by 2-dimensional-gel electrophoresis followed by western blot using an anti-YB1 antibody. As a loading control, ß-actin was detected. The pI of unacetylated YB-1 is pH5.74. (C and D) Autophagy inducing activity was compared between iDC and 27DC using the Cyto-ID reagent as previously described [2]. iDC and 27DC were incubated 6 hours in a 96-well plate. Autophagosome staining was performed using the Cyto-ID reagent and Hoechst 33342 as a counterstain. DAPI staining was conducted as a nuclear counterstain, (C) 2×3 composite images were taken on a Zeiss AxioObserver motorized microscope. (D) Autophagy induction in iDC and 27DC was measured. The average stained area per cell for each condition was calculated [24]. Data shows means±SE (n=4).

Autophagy assay demonstrated that endogenous autophagy induction in 27DC was not significantly changed (*p*=0.063, n=5) (**Figure 4C**, **4D**), which differs from IL-27-induced macrophages [24]. Based on these data, we concluded that 27DC suppressed RT step using a mechanism different from the ones we previously reported in MDMs and T cells.

### Comparison of gene expression profiles

To define the host factor(s) associated with HIV resistance in 27DC, we compared the gene expression profiles between iDC and 27DC with DNA microarrays. The profiles were compared among four donors’ iDC and 27DC **(Figure 5A).**

**Figure 5.**
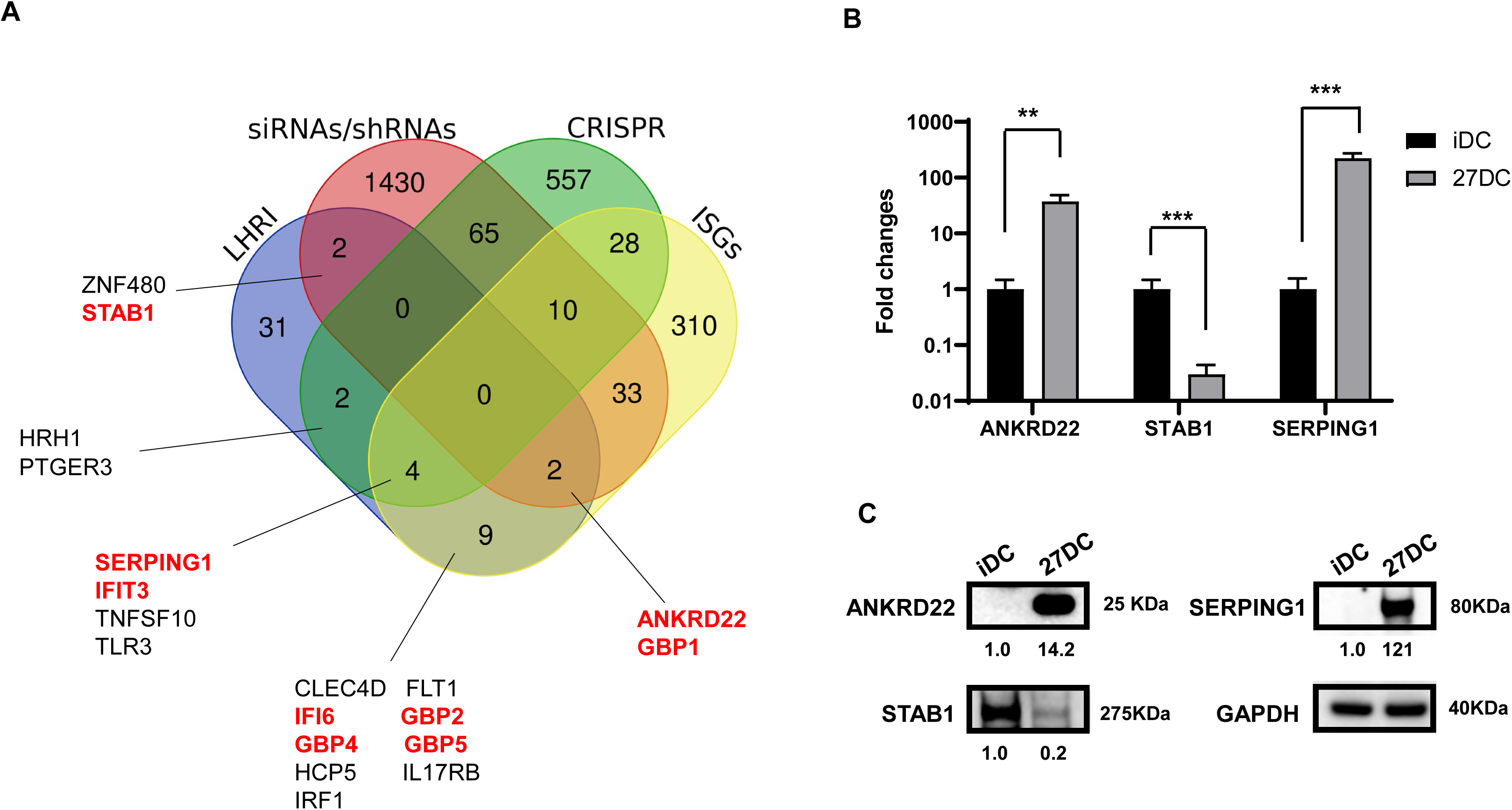
Comparison of gene expression between iDC and 27DC. (A). A microarray analysis was performed on iDC and 27DC from 4 different donors, and a total of 51 genes (**Supplemental Table 1)** were significantly changed by > 3-fold in 27DC (*p*<0.05). The Venn diagram analysis was conducted using reported genes associated with HIV replication by siRNA/shRNA screening (siRNAs/shRNAs), CRISPR libraries screening (CRISPR), and the reported Interferon-stimulated genes (ISGs). Of 51, 19 genes overlapped with the reported genes. The gene names with the red font were targeted for further analysis. The genes with black font encode receptors or transcriptional factors. (B) The qRT-PCR was conducted to define the expression level of each gene of interest using total cellular RNA from iDC and 27DC from three independent donors. Data show mean ± SE from three independent donors’ cells. (C) The total cell lysate was collected from iDC and 27DC, and Western blotting was performed using anti-ANKRD22, anti-STAB1, anti-SERPING1, and anti-GAPDH antibodies. The band intensity of each protein was normalized by the band intensity of GAPDH (Image J); the values are indicated below images.

A total of 51 genes, commonly either up- or downregulated in four donors by more than 3-fold (*p*< 0.05) (**Supplemental Table S11),** were cross-referenced with 2,452 previously reported host factors associated with HIV replication **(Supplemental Table S1).** These factors were identified during screening of siRNA, shRNA libraries [65–69] or CRISPR libraries [70–79] and potent IFN-stimulated genes (ISGs) [80]. Cross-reference analysis demonstrated that 19 genes were hit as potentially associated with HIV resistance in 27DC. As mentioned above, 27DC suppressed RT step after infection, thus, total 10 genes encoding transcriptional factors, or cell surface receptors were excluded from the common 19 genes. Among the remaining 9 genes, anti-viral effects of GBP1, GBP2, GBP4, GBP5, IFIT3, and IFI6 have been reported. GBP2 and GBP5 but not GBP1 and GBP4 suppress HIV infection. Those proteins are incorporated in virus particles and suppress HIV infection [81, 82]. As shown in Figure 2A, HIV derived from 27DC had a comparable infectability to the virus from the iDC, indicating even though GBP2 and GBP5 were induced in 27DC, the proteins do not play a key role in the anti-HIV effect in 27DC. IFIT family proteins, including IFIT3 are induced after virus infection [83, 84] and suppresses translation of viral proteins [84]. IFI6 is reported as an anti-HIV proteins in HeLa [85] but not in Jurkat T cell [86]. The anti-HIV effect of IFI6 in Hela cells is caused by the autophagy induction. As shown in Figure 4C, 27DC did not induce autophagy; thus, it appears that IFI6 does not function as an anti-HIV in 27DC. Therefore, IFIT3 and IFI6 were also excluded from the potent anti-HIV factor(s) in 27DC. Although Stabilin 1 (STAB1) is a cell surface receptor [87, 88], it is known that the STAB1 expression regulates the production of CCL3, an R5 HIV binding inhibitor [89] and as shown in Figure 2A, HIV binding was inhibited on 27DC; therefore, we included it in the downstream analysis. Therefore, remaining three genes: ankyrin repeat domain 22 (ANKRD22), serpin family G member 1 (SERPING1), and STAB1 were further analyzed. To confirm the expression of each gene, qRT-PCR was performed. The gene expression levels in 27DC were normalized to the expression of an internal control, GAPDH, and compared to the expression levels in iDC **(Figure 5B).**

The expression of each protein was confirmed by Western blotting using whole-cell lysates of IDC and 27DC **(Figure.5C).** The expression of ANKRD22 and SERPING1 in 27DC increased by 14- and 121-fold, respectively, compared to that in iDC. In contrast, STAB1 expression in 27DC was suppressed by 80% compared to iDC.

### Evaluation of a role of ANKRD22 expression in 27DC on HIV replication

ANKRD22 is a member of the ankyrin family [90–92], and the ankyrin protein is considered a novel viral entry inhibitor [93]. It is reported that the expression of ANKRD22 is associated with chronic HIV infection in macrophages and T cells [94]; however, the impact of ANKRD22 on HIV replication, focusing on its role in the RT reactions, has not been reported. As a result, we attempted to evaluate the ANKRD22 effect on RT reaction by knocking down ANKRD22 using siRNA against ANKRD22 (si-ANKRD22). To suppress the inducing ANKRD22 mRNA, we first performed a kinetic study of its expression. Monocytes were cultured in the absence or presence of IL-27, and cells were harvested from day-1 to -7. We found that ANKRD22 was induced in IL-27-treated cells within 24 hours **(Figure 6A),** thus we decided to transfect si-ANKRD22 into monocytes before starting IL-27 treatment.

**Figure 6.**
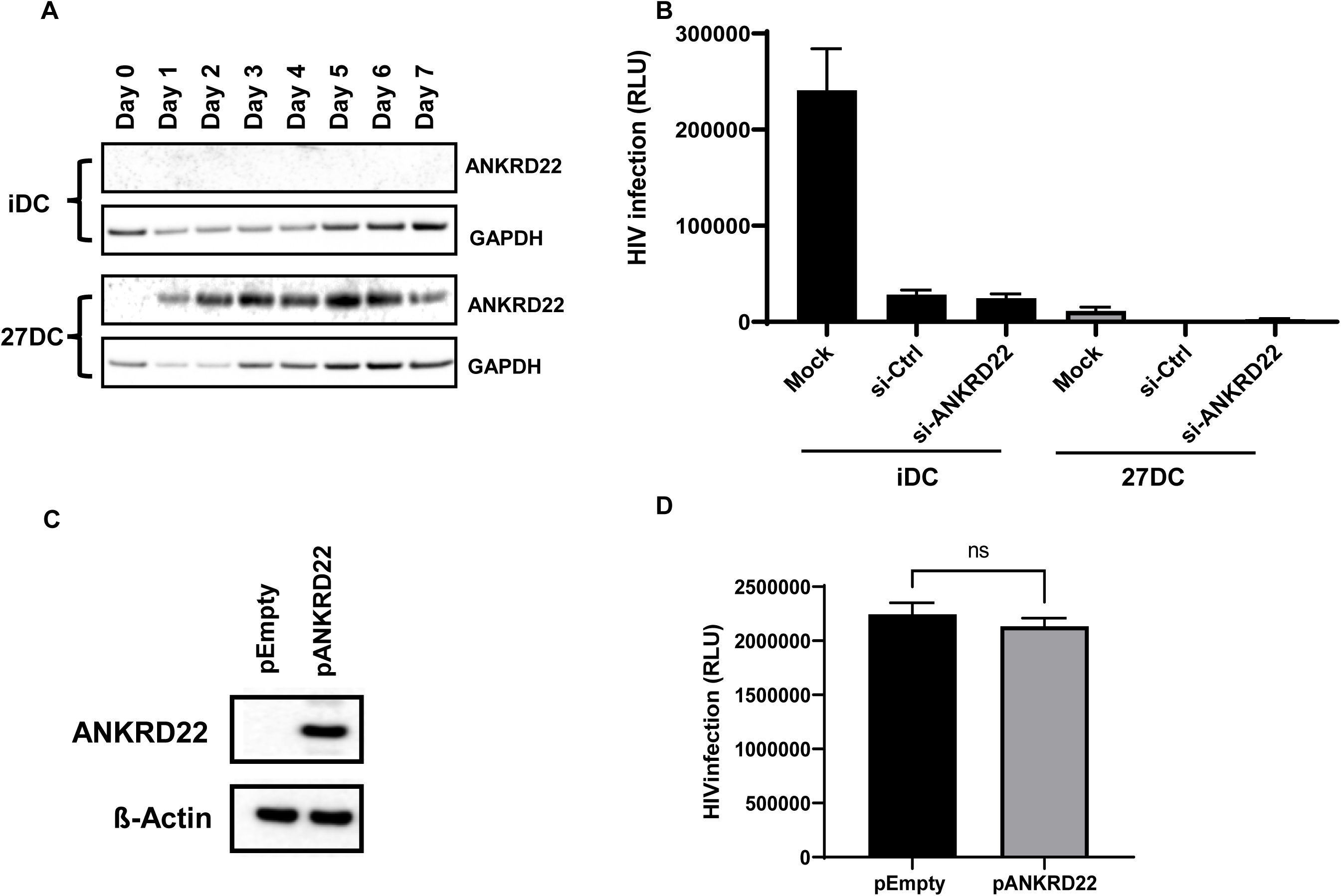
ANKRD22 expression has no impact on HIV infection. (A) Monocytes were differentiated into iDC or 27DC in G4 media in the absence or presence of IL-27. Cell aliquots were collected every day and whole cell lysate was subjected for detecting ANKRD 22 expression by WB using anti-ANKRD22 antibody. As internal loading control, GAPDH expression was detected. (B) Control siRNA (si-Ctrl) and siRNA targeting ANKRD22 (si-ANKRD22) (Thermo Fisher) was transfected into monocyte using HiPerFect, and then differentiated to iDC or 27DC. The cells were recovered and then infected with HIVLuc as described in the Materials and Methods, Luciferase activity on 48 hours after infection was quantified. A representative result from three independent assays is presented as means ± SDs (n = 5). (C, D) empty plasmid (pEmpty) or ANKRD22-encoding plasmid (pANKRD22) was transfected into HEK293T cells for 48 hours, aliquots of cells were subject for Western blot to detect ANKRD 22 expression (C) and remaining cells were infected with HIV-Luc (D) and then HIV infection was quantified by Luciferase activity. Representative data from three independent assays are presented as means ± SDs (n = 3).

The si-ANKRD22 and a control none-targeting siRNA (si-Ctrl) were delivered using several different transfection lipids: Lipofectamine RNAiMAX (Thermo Fisher), Lipofectamin2000 (Thermo Fisher), TransIT-siQUEST (Mirus), INTERFERin (Polypus), or HiPerFect T (Qiagen), and electroporation method using the 4D system with the P3 reagent (Lonza). Even though we used different sequences of si-ANKRD22 (obtained from Thermo Fisher, Horizon, or OriGene), none of si-ANKRD22 consistently suppressed ANKRD22 expression in 27DC (data not shown). However, as shown in **Figure 6B**, si-ANKRD22 administration did not restore HIV infection in 27DC but suppressed HIV infection in iDC by 89.7 ± 4.1% (n = 5). Of note, the transfection of si-Ctrl and si-ANKRD22 significantly suppressed HIV infection in iDC by 88.2±4.4 % and 89.7±4.0 %, respectively, suggesting that an off-target effect by siRNA transfection suppresses HIV in DCs.

Transfection of siRNA induces the innate immune response and produces type I and III IFNs as one of off-target effects [95, 96]; the induced innate immune responses may suppress HIV infection. Accordingly, we could not define a role of ANKRD22 using siRNAs. We were, therefore, not able to define a role of ANKRD22 expression in 27DC using siRNAs.

As an alternative approach, the overexpression of ANKRD22 in iDC was attempted. A plasmid DNA encoding ANKRD22 gene (pANKRD22) was transfected into iDC or monocytes using lipids: Lipofectamin2000 (ThermoFisher), JetPEI (Polyplus), FuGENE6 (Promega), TransIT-LT1 (Mirus Bio), and Efffecten (Qiagen), or electroporation using P3 reagent; however, all attempted methods displayed strong cytotoxicity, thus we failed to define a role of the overexpressed ANKRD22 on HIV infection in iDC. To define a role of ANKRD22 expression, we transfected the pANKRD22 into HEK293T cells, which do not induce IFN production in response to transfected DNA because of the lack of STING [97]. Although transfection induced the gene product (**Figure 6C**), the expression had no significant impact on HIV infection using pseudotyped HIVLuc-VSVG (**Figure 6D**).

### Evaluation of a role of STAB1 or SERPING1 in 27DC on HIV replication

STAB1 is a cell surface protein and acts as a scavenger receptor [87, 88]. It is reported that the suppression of STAB1 expression is associated with increased production of CCL3 (also known as macrophage inflammatory protein (MIP)-1 alpha) without an increase in CCL3 gene activation [89]. Although the mechanism of the STAB1-mediated regulation of CCL3 production is unknown, If the 27DC produces CCL3 due to the down regulation of STAB1, the induced CCL3 might affect the suppression of HIV binding, thus, we assessed the secretion of CCL3 during the induction of 27DC. Culture supernatants from iDC and 27DC were collected on day 7, and the CCL3 was quantified. The results indicated that CCL3 was significantly secreted in 27DC culture supernatants **(Figure 7A).** Microarray analysis and qRT-PCR assay demonstrated that compared to iDC, CCL3 gene expression was changed in 27DC by 1.37 ± 0.28-fold (n = 4) and 2.2 ± 0.06-fold (n = 3, p=0.317), respectively. These results consisted of previous reports [89]; the downregulation of STAB1 induced CCL3 protein without a change in the gene expression. CCL3 is a member of CCR5 ligands; therefore, we also examined the secretion of other CCR5 ligands, CCL4 (MIP1-beta) and CCL5 (RANTES), in the culture supernatants. The quantification assay displayed that CCL4 expression was not significantly changed and CCL5 protein was not detected (below detection limits) in 27DC culture supernatants (Figure 7 A). The qRT-PCR demonstrated that CCL4 expression in 27DC was not significantly changed (2.4 ± 0.4- fold, p=0.17, n=3) (Figure 7B); on the contrary, CCL5 expression was 19.6 ± 8.7-fold increased (p<0.01, n=3) in 27DC, compared to iDC. Overall, 27DC uniquely produces CCL3. This inducing activity in 27DC may contribute to the partial suppression of HIV binding (Figure 2A).

**Figure 7.**
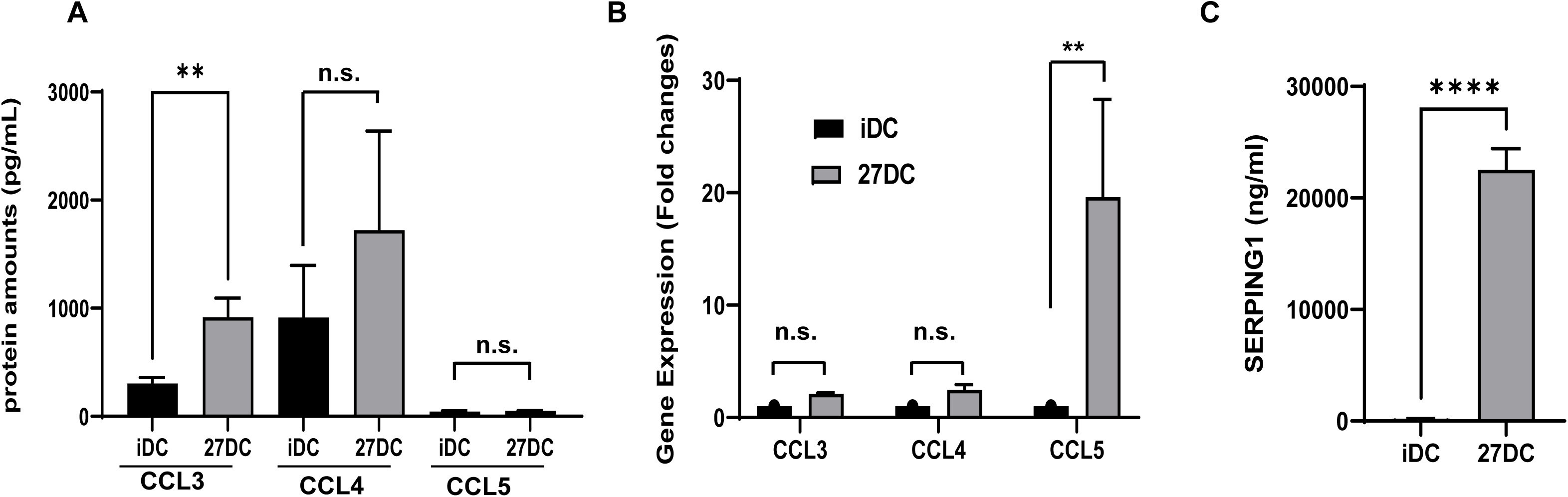
27DC produce CCL3 and SERPING1 in culture supernatants. (A) The concentrations of secreted CCL3, CCL4 and CCL5 in culture supernatants on day 7 were measured by ELISA. A total of eight different donor culture supernatants were subjected to the assay. (B) Real-time RT-PCR was conducted to compare the expression level of CCL3, CCL4 and CCL5 gene. Data show mean ± SE from three independent donors’ cells. (C) The concentrations of secreted SERPING1 in culture supernatants on day 7 were measured by ELISA. A total of eight different donor culture supernatants were subjected to the assay. Data indicates means ± SE (n=8). **: *p*<0.01, and ***: *p*<0.001.

SERPING1, also known as C1 inhibitor (C1INH), is a member of the plasma serine protease serpin superfamily [98]. It is mainly produced from the endothelium and inhibits both the classical (C1r and C1s) and mannose-binding lectin-associated serine protease (MASP)1/MASP2 complement pathways, as well as enzymes involved in fibrinolytic pathway, intrinsic coagulation, and contact systems, such as plasmin, factor XI, factor XII, and plasma kallikrein [98, 99]. It is induced by IFNs [100, 101]. To define whether SERPING1 is produced from 27DCthe protein amounts was quantified using an ELISA kit, The results indicated that SERPING1 was significantly produced from 27DC **(Figure 7C),** suggesting that IL-27 can induce the protein like IFNs, and it may affect HIV infection during differentiation. To assess anti-HIV effect of SERPING1, iDC was cultured with 20 µg/mL of recombinant SERPING1 proteins for three days and then HIV infection assay was conducted using HIVLuc-VSVG. The result displayed that SERPING1 had no impact on HIV infection (**Supplemental Figure 1**)

### Conclusions

The current study revealed that IL-27 differentiated monocytes into HIV-resistant DCs, suppressing R5 HIV binding to the cells and the HIV RT reaction after viral entry. This suppression mechanism differs from the previously reported IL-27-induced HIV-resistant macrophages or T cells; the anti-viral effect mediated via SPTBN1, autophagy, or YB1-dependent mechanisms [24, 42, 58], indicating that IL-27 can induce HIV-resistant cells in a cell type-dependent manner. Due to failing knockdown by siRNA or overexpression of protein by plasmid transfection in 27DC or iDC, the molecular mechanisms underlying the inhibition of HIV in 27DC remain unknown. We found that 27DC induces CCL3 and SERPING1 production during differentiation; CCL3 protects against R5 HIV infection in DC, macrophages, and T cells [102]; therefore, if 27DC is cultured with other cell types, macrophages or T cells, the secreted CCL3 may inhibit HIV infection in the cocultured cells by *trans*.

SERPING1 functions as the major inhibitor of C1r, C1s, mannose-binding lectin-associated serine protease MASP-1, MASP-2, factor XII and kallikrein in the contact system, factor XI and thrombin in the coagulation system, and tissue plasminogen-activator and plasmin in the fibrinolytic system [103–106] and SPERPING1 is proposed as a useful adjunct in the management of COVID-19 with severe pneumonia [107]. Thus, IL-27 primarily affects HIV-1 replication and potentially other viral infections, including SARS-Cov2 infection. Further research is needed to provide new insights into biological function of IL-27 and cellular function of 27DC and explore IL-27’s potential as a therapeutic cytokine.

## Acknowledgments

Authors thank HC. Lane and MW. Baseler for supporting this project, W. Greene for kindly providing pHIV-1_NL4/3_-Env_AD.8_-ΔVpr and pVpr-BLAM, D. Poudyal for discussion, M. Bosche for technical support. All authors thank MA. Martin for providing pNL4.3 and pAd8, and D. Yang for providing reagents. Authors thank W. Chang for critical reading. This project has been funded in whole or in part with federal funds from the National Cancer Institute, National Institutes of Health, under Contract No. HHSN261200800001E. The content of this publication does not necessarily reflect the views or policies of the Department of Health and Human Services, nor does mention of trade names, commercial products, or organizations imply endorsement by the U.S. Government. This research was supported [in part] by the National Institute of Allergy and Infectious Disease.

## Author Contributions

TI created the project, designed experiments, performed assays, and wrote the manuscript. QC and BS designed experiments, performed assays. SL designed and performed assay. and TI, QC, JY, SL, JA, JH, and HS analyzed the data. BS, JY and SL wrote methods. AM and CW performed assays. All authors reviewed and approved the final version of the manuscript.

---

**Supplementary Table S1.**
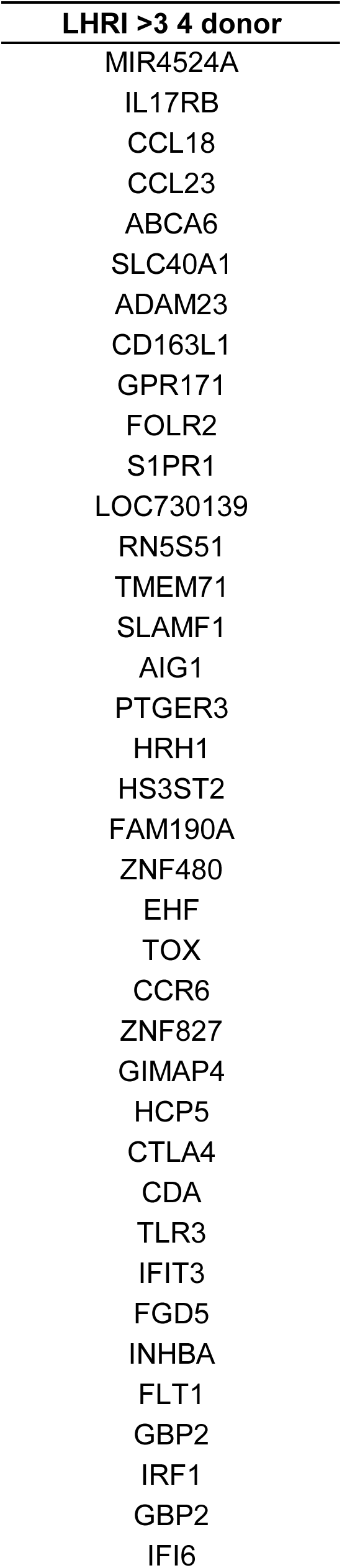
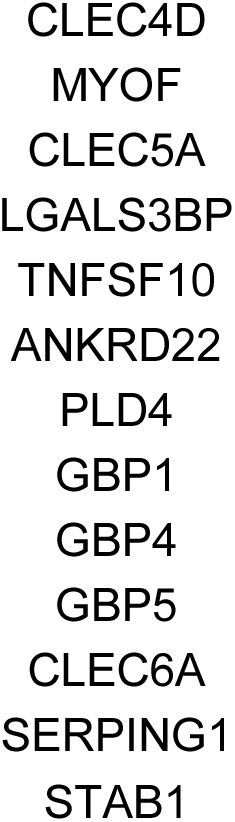
Gene list commonry changed 3 fold in 27DC (p<0.05)

**Supplemental Figure S1.**
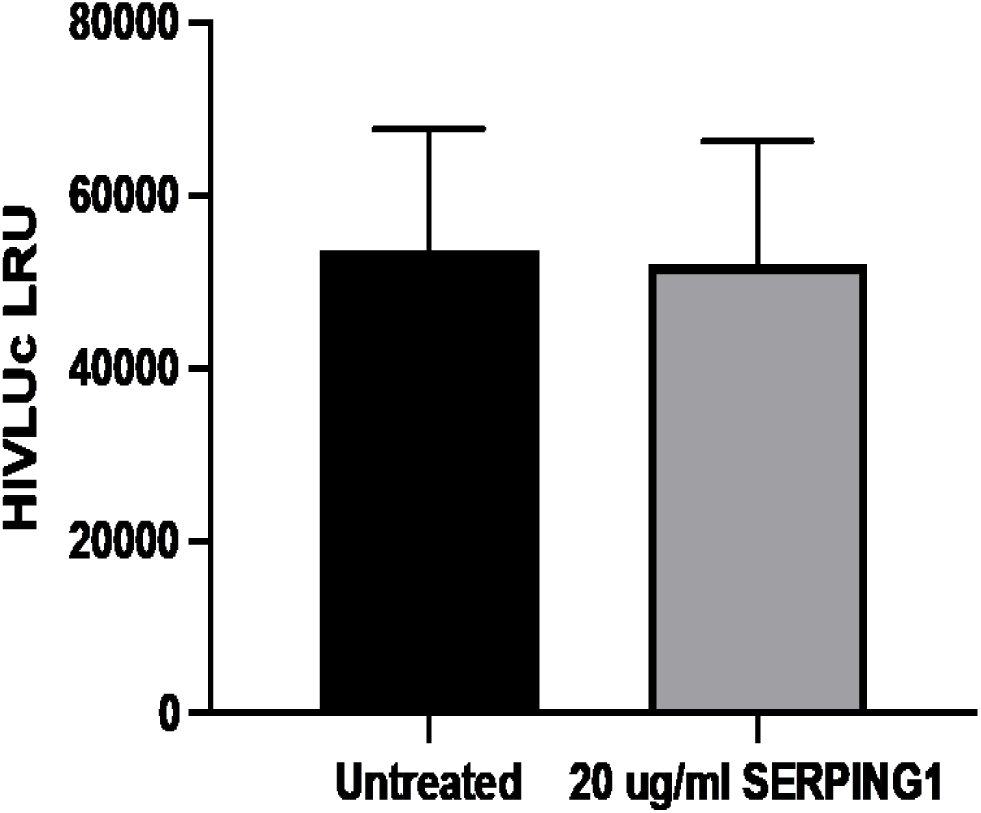
iDC were cultured with 20 µg/mL of recombinant SERPING1 for 3 days and then infected with HIVLuc as described in the Materials and Methods. HIV infection was quantified by luciferase assay. Data show Mean ± SD (n=3) of a representative data from two independent assays.

